# Intent matters: how flow & forms of information impact collective navigation

**DOI:** 10.1101/2023.06.23.545920

**Authors:** T. M. Hodgson, S. T. Johnston, M. Ottobre, K. J. Painter

## Abstract

The phenomenon of collective navigation has received considerable interest in recent years. A common line of thinking, backed by theoretical studies, is that collective navigation can improve navigation efficiency through the ‘many-wrongs’ principle, whereby individual error is reduced by comparing the headings of neighbours. When navigation takes place in a flowing environment, each individual’s trajectory is influenced by drift. Consequently, a potential discrepancy emerges between an individual’s intended heading and its actual heading. In this study we develop a theoretical model to explore whether collective navigation benefits are altered according to the form of heading information transmitted between neighbours. Navigation based on each individual’s intended heading is found to confer robust advantages across a wide spectrum of flows, via both a marked improvement in migration times and a capacity for a group to overcome flows unnavigable by solitary individuals. Navigation based on individual’s actual headings is far less effective, only offering an improvement under highly favourable currents. For many currents, sharing actual heading information can even lead to journey times that exceed those of individual navigators.

## 1 Introduction

Migration occurs in all major animal groups, from arthropods to birds; and can occur across vast distances. Frequently, migration is a synchronised movement, with as few as a pair to a vast number travelling *en masse* [1]. For those animals able to sense the presence of others, this provides a potential source of navigating information.

Long distance migrations require overcoming multiple challenges, including energy management, avoiding predation, coping with adverse environmental conditions, and finding the destination site [2]. Robust and efficient navigation can potentially mitigate the impact of these challenges. Accordingly, questions arise regarding the mechanisms that are used to reach a destination in a relatively short time. Such questions, and animal migration in a wider sense, have fascinated scientists for centuries, as far back as Aristotle [3]. Since the advent of mark-and-recapture studies, and more recently radar and tracking devices, our knowledge of animal movement has expanded greatly and new mechanisms of navigation have been proposed (see for example [4] and references therein).

When navigating within a group, an individual can be considered to have two sources of information: *inherent information* and *group information*. The former refers to the navigation information an individual has direct access to, for example by sensing environmental cues (odour plumes, stars, magnetic field etc.) or accessing stored or inherited information (memory). It can also be viewed as the individual level of confidence concerning the direction to the goal. By group information we refer to the information gained by observing or receiving the headings of other individuals. A conventional line of thinking is that group information can provide navigational benefit through the many-wrongs principle. Individual errors are suppressed by averaging across the group [6, 7]. Group cohesion rules, present in many collective movement models, can also confer indirect navigational benefits via encouraging individuals to aggregate [8]. A number of modelling studies have tested this thinking, corroborating the benefits of the many-wrongs principle, albeit for simpler navigational scenarios [9, 10, 11, 12].

Dynamic fluid environments present a particular challenge to navigation. Individuals and populations can be heavily impacted by passive advection (ocean currents, prevailing winds etc.) and a robust navigation strategy may be necessary for recovery of direction in the event of significant disturbances. However, the subtleties of just how flow impacts on collective migration have received relatively little attention to date [12]. In a flowing environment, the direction an individual intends to travel and the direction they actually travel may not coincide [13, 14]. This could have repercussions on collective navigation benefits, depending on the type of information communicated. It also raises the question of whether individuals compensate for the effect of the flow in their headings, as shown experimentally in certain avian species [15, 16].

Many animals have an ability to sense their environment and detect the presence of others. A (significant) subset of those, such as whales, also have the ability to send signals that can be received by others: an ability to communicate. The ‘language’ of this communication has been studied in many species [17], with the ‘singing’ of species such as the humpback whale (*Megaptera novaeangliae*) in particular capturing popular attention [18, 19, 20].^1^ While the physical transmission of these signals is relatively well understood, the question of what information can be communicated and be received remains largely unsolved at the level of a particular species. Do members only respond to simple observations of other individuals (i.e., a passive signal)? For example, do they align directions according to those of neighbours, as proposed for starlings [21]? Or do they respond according to information that is actively communicated from another individual? Can an individual communicate its intended direction? If so, what impact does this have on navigation success at a population level?

Modelling studies can provide insight into these (and other) questions, and can be especially beneficial for systems in which there is a difficulty of obtaining data. Many models for collective behaviour exist (see [11] for a general review), covering a broad range of intricacy. As far as we are aware, none of these have explicitly incorporated a turbulent environment and the different ways that this could impact on communication. Precisely determining the type of information transmitted by a species is almost impossible with current methods [22], and modelling therefore provides a technique through which to test different degrees of communication and their impact on animal navigation.

In this study, we perform a computational modelling study into collective migration in a flowing environment. Section 2 contains an introduction to the model. In Section 3 we describe our results, systematically exploring collective navigation success across a range of simple and complex flows. In Section 4 we discuss the results and implications in the context of real-world navigation.

## 2 Models and Methods

We describe the movement of an individual via a velocity-jump random walk (VJRW). [23, 24, 25]. In a VJRW, an individual moves with constant velocity in a straight line for a random length of time before randomly selecting a new velocity. The times at which tumbles occur is governed by a time-homogeneous Poisson process with rate *λ*, which we refer to as the *turning rate.* Similar models are common in the study of collective behaviour, see for example [10]. In the formulation described below, we consider alignment of velocities (as [26] etc.) but do not incorporate an explicit attraction or cohesion (as [27, 28] etc.). The key novelties lie in the inclusion of turbulent background flows and the heading information that is communicated. Note that for the present study we do not include any drift compensation, as has been described in many species, particularly birds [29, 15, 16, 14]. However, in Section 4 we describe preliminary results for a model that includes a form of drift compensation, and provide an expanded discussion that highlights the considerable number of subtleties to be accounted for when incorporating this into the framework.

Suppose we have a population of *N* individuals, where the *i*th individual has position *x_i_*(*t*) ∈ ℝ^2^ at time *t*. Each individual’s position is governed by

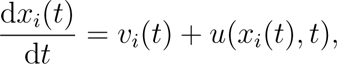

where *v_i_*(*t*) ∈ ℝ^2^ is the *active* velocity of the individual: the component of movement due to self-generated propulsion. The component of the velocity caused by the surrounding medium is given by *u*(*x_i_*(*t*)*, t*), which we refer to as the *passive* velocity. We assume, for simplicity, that individuals move in two dimensions, for example at the surface of the ocean. For convenience, we will assume the active speed is fixed, i.e., *|v*(*t*)*|* = *s* for every *t ≥* 0. For a general setting it is convenient to assume an *a priori* scaling of length and time scales such that *s* = *λ* = 1. This scaling can, of course, be reversed in the context of a particular application. Note that we neglect hydrodynamic interactions between the individuals’ bodies and the fluid, assuming that at the scales of interest the individual can be regarded as a point mass.

### 2.1 Navigation

We assume that the principal objective of each individual is to reach a target destination, centred at position *x^⋆^ ∈* ℝ^2^, which we refer to as the *goal*. To model this navigation process, we assume that at each reorientation event an individual selects a new heading drawn from a probability distribution that combines both inherent and group information. The inherent (and group) information is encoded as a von Mises distribution. In Figure 1c) we show an example of a constant information field, the inherent information distribution from that field, the group information distribution and the combination into a heading distribution. The next section gives details on the construction of these distributions and the combination into a heading distribution.

**Figure 1:**
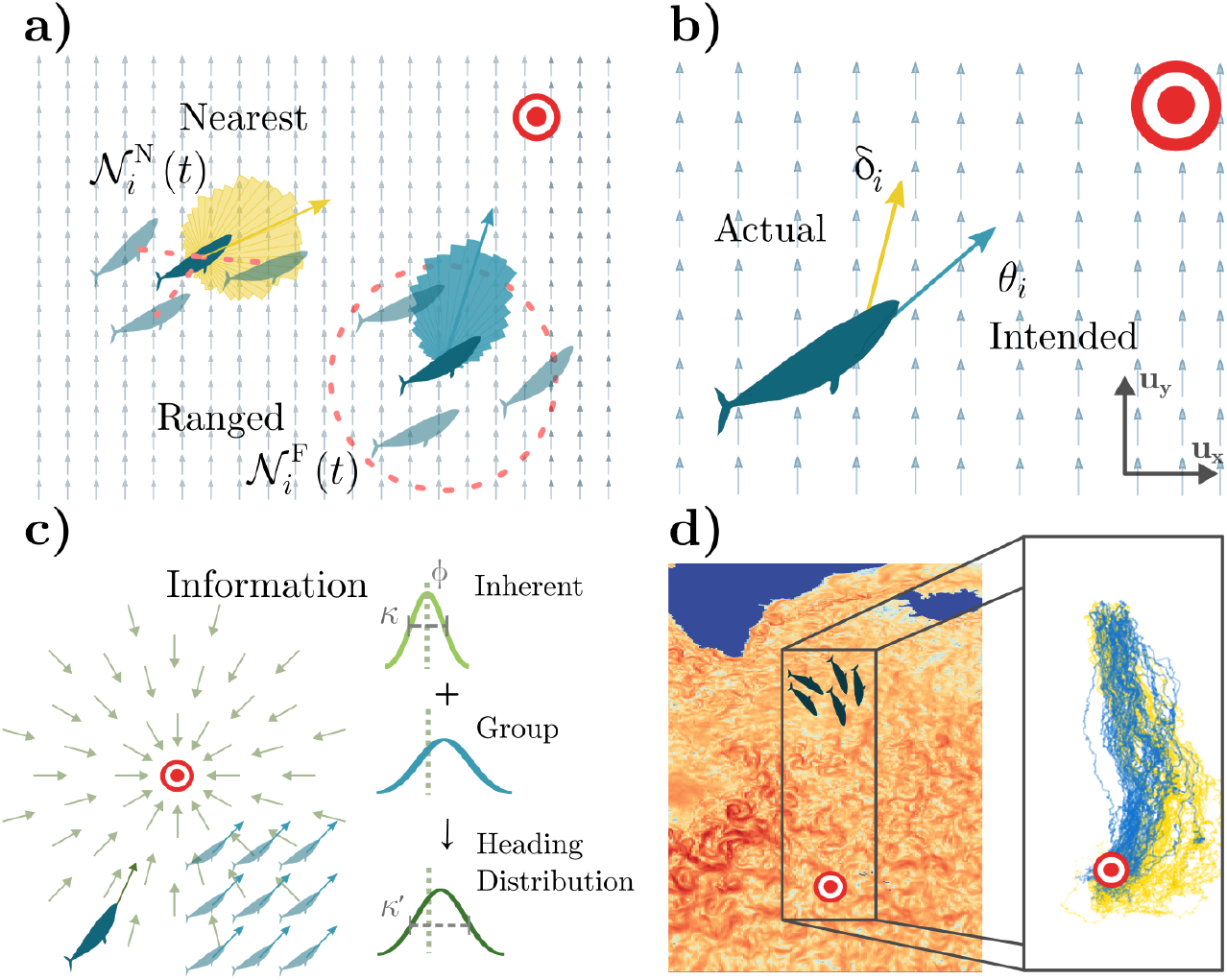
Model Summary. a) Interacting with nearest neighbours 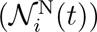 as opposed to a fixed radius interaction, 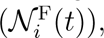 changes the potential heading distributions (coloured rose plots). Interaction is shown using a red dashed line. b) An individual’s intended heading *θ_i_* (blue) can differ from their actual heading *δ_i_* (yellow) in a flowing environment when navigating towards a target (red/white). Light blue arrows indicate the background flow direction. c) Information is gathered from the environment and neighbours to form a heading distribution for sampling during reorientation. Light green lines indicate the goal direction. d) Ocean flow data from the North Atlantic and simulation region used in Section 3.3 with example trajectories navigating using the intended (blue) and actual (yellow) headings [5].

#### 2.1.1 Inherent Information

In this setting, inherent information represents the confidence an individual has in the position of the goal relative to its current position. We model the inherent information as a vector field (see Figure 1c), in which the direction and magnitude of a vector at a position *x_i_* indicates the direction towards the goal and the confidence in the knowledge, respectively. Specifically, we assume the goal direction is encoded in a location parameter, *ϕ*, while the confidence is encoded in a concentration parameter, *κ*, of a von Mises distribution with density *f* given by [30]:

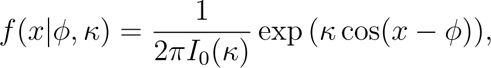

where *I*_0_(*κ*) is the modified Bessel function of the first kind of order zero.^2^ We denote the sampled heading from this von Mises distribution *θ^′^ ∼ vM* (*ϕ, κ*). If the *i*th individual is at position *x_i_*(*t*), then the direction towards the goal is described by the angle

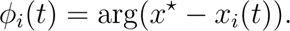

Larger *κ* corresponds to increasing confidence in the goal direction. For the present study our main results are presented for *κ* = 1. For context, this means an individual will reorient within 45*^◦^* of the goal direction approximately 50% of the time and is chosen to ensure a balance between overall navigation to the goal and a non-negligible uncertainty in the available information. A sensitivity analysis is performed for different constant values of *κ* in Supplementary Information C. The work of [31] considers the scenario in which the level of information (and thus *κ*) varies with position in a non-flowing environment.

#### 2.1.2 Group Information

To include interactions within a group, the reorientation mechanism also depends on the locations and headings of neighbouring individuals. There are three key points to consider: what defines a neighbour, what information is communicated, and how the individual combines that information with its inherent information.

##### Neighbour Selection

Given a certain sensing mechanism, *d*, we define the set of neighbours of individual *i* at time *t* as 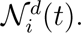 We consider two sensing mechanisms: *ranged* (fixed radius) interaction (F) and *nearest neighbour* interaction (N), such that *d ∈ {*F, N*}*. Correspondingly, we will consider the sets 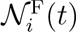 and 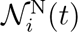 defined as below.

- For the ranged interaction, we assume that individuals have a fixed sensing range *R ∈* ℝ^+^ and that all individuals within *R* can be detected. Subsequently we consider

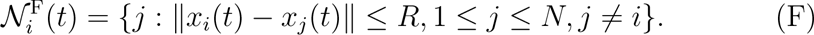 Here ‖ · ‖ denotes the Euclidean 2-norm. Note that our model assumes that all neighbours within 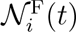 are considered equally. Ranged interaction may describe the finite propagation of a signal through a medium, or the limit to a species’ auditory or visual ability.
- In the nearest neighbour interaction, an individual considers only the *S* closest individuals to itself while reorienting, irrespective of the distance between them. That is,

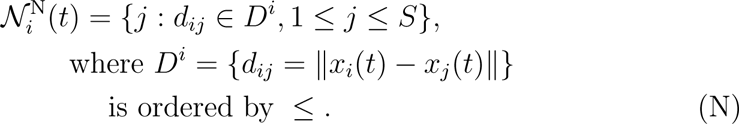 This was proposed as a possible model within starling flocks in [21]. Nearest-neighbour sensing could feasibly account for the maximum cognitive load an animal is capable of sustaining.

Of course, it is probable that for many species a combination of these two may operate: an individual will be limited by the distance and the cognitive load it can process. By selecting these two generic models, our aim is to test robustness with respect to two distinct methods for selecting neighbours. Figure 1a) provides an illustration of these two methods of neighbour selection.

##### Communicated Information

Given these two methods of neighbour detection, we must next consider what information can be transferred between navigating individuals. In the absence of flow, an individual simply moves in the direction *θ_i_* due to active movement, which we call the *intended heading*. In a flowing environment, however, the actual heading may be distinct from the intended heading due to drift. This leads us to investigate two forms of information transfer, according to whether an individual receives (or detects) intended or actual headings.^3^ The difference in these heading and perception types is illustrated in Figure 1b).

- The intended heading, (I), is the active swim direction of the individual *i*,

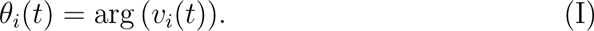
- The actual heading, (A), is the flow-affected direction of the individual *i*,

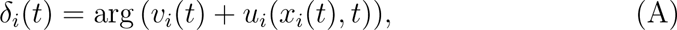 i.e. the combination of the passive and active movement.

##### Combining Group and Inherent Information

We now describe how the group information is combined and integrated with the inherent information to guide reorientation decisions. First, a resultant vector that combines the circular mean of the neighbour headings and the newly sampled heading based on the individual’s inherent information is calculated, *θ^′^*. That is, the resultant vector for a navigator *j* at time *t* is

Intended:

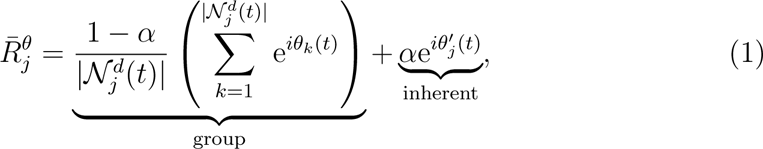

Actual:

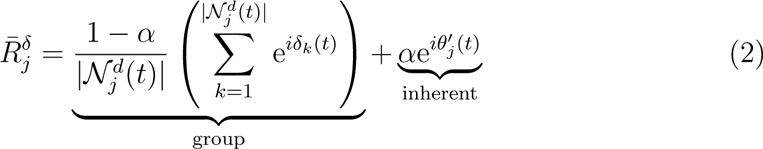

where 0 *≤ α ≤* 1. The argument of the resultant vectors are denoted

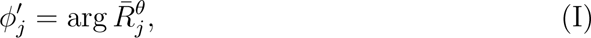

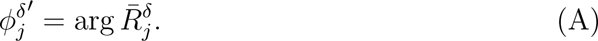

The parameter *α* is a weighting between the headings based on the group and inherent information. Here we use an equal weighting (*α* = 1*/*2) which, following a sensitivity analysis, was shown to broadly give the fastest successful migration times under a variety of information fields [31]. The combination is the same irrespective of the heading type (actual, (A); intended, (I)) and sensing mechanism (nearest-neighbour, (N); fixed-ranged, (F)). To capture the uncertainty in the model, as expected in a noisy biological environment, the calculated heading 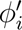 is instead used as the mean of a von Mises distribution. Specifically, the new heading for the reorienting individual is drawn from

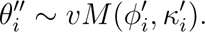

The circular mean described above is the maximum likelihood estimator for the mean of a von Mises distribution. To choose an appropriate value for the concentration parameter, *κ^′^*, given the number of neighbours, we preferentially use a pre-calculated lookup table of the likelihood. This method is expanded upon in [31], we only note here that it avoids biases in the estimates of *κ^′^*, calculated by repeatedly generating samples for fixed *|N |* and *κ* and determining the uncorrected *κ^′^* value for each sample. Algorithms for this method are given in the Supplementary Information. Given these mechanisms, we now have 4 distinct submodels, as summarised in Table 1.

**Table 1:**
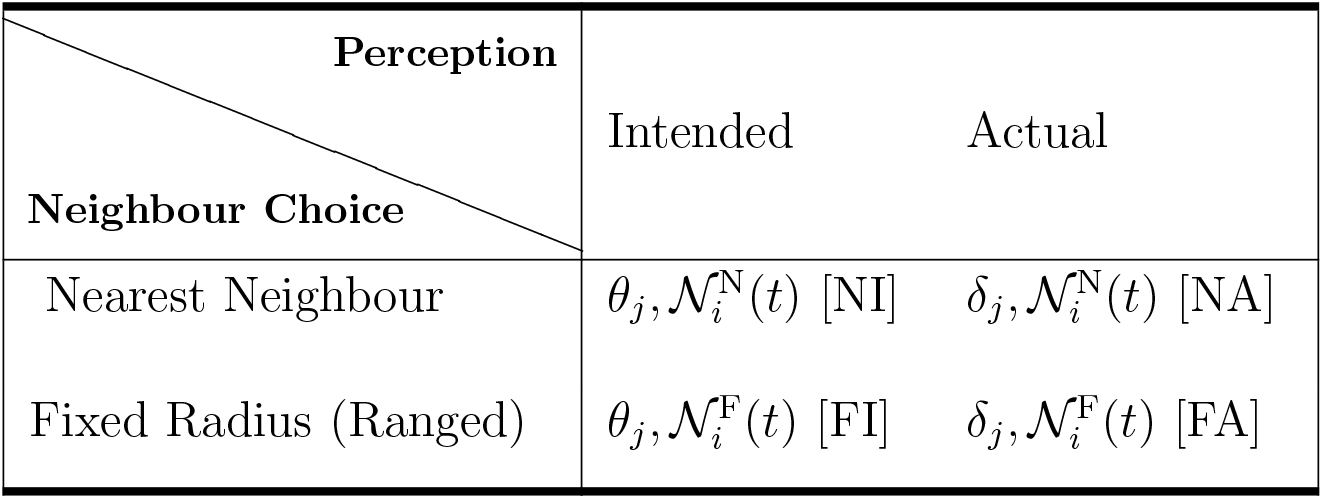
Summary of the models used. Recall *θ_j_* is the intended heading direction, and *δ_j_* is the actual heading direction. 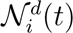 is the set of neighbours of the *i*th individual at time *t*, using distance metric *d*. The model abbreviations used in the body text are given in square brackets.

### 2.2 Goal Location and Initial Distribution

An individual is defined as arrived at the goal when it is within a distance *ε* of the target position *x^⋆^*:= (0, 0). That is, the individual has arrived if

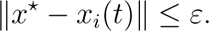

The parameter *ε* is referred to as the *goal tolerance*. Once an individual has arrived, it is removed from the simulation and has no further impact on individuals that are still navigating. In the abstract setting here, this is a modelling choice and we discuss this further in Section 4.

The initial positions of the individuals for all simulations are drawn uniformly at random from a square. More specifically,

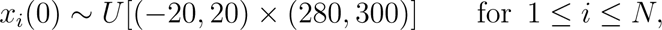

where *U* [*·*] denotes the uniform distribution on the set.

Note that an individual is said to have *failed* if it does not reach the goal before a terminal time *T* = 5000, or if its distance from the goal at any time exceeds 1.5 times its initial distance from the goal. The group is said to have failed if more than 50% of individuals fail. These values are primarily dictated by computational considerations, however we note that for the initial position considered this is a value more than 15 times larger than the shortest possible time taken for an individual to arrive. To measure group success, we consider the median arrival time as it is a fairly general measure that is relatively robust to parameter variation – see Supplementary Information B, where we also implement the *navigational efficiency* as considered in [32, 28, 8].

### 2.3 Background Flow Fields

We consider two forms of environmental flow: constant (in space and time) flow and quasi-real-world flow scenarios. The first is described by a flow angle *ϑ* and flow strength *ζ ≥* 0 such that

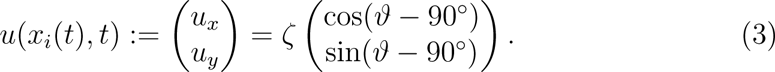

Note that due to the normalisation of each individual’s active movement speed, a flow strength of *ζ* = 0.25, say, would indicate that the flow is 25% of the individual’s movement speed. Given the above initial positions and target location, if *ϑ* = 180*^◦^*, the flow is opposing the direction to the goal and we call this an *unfavourable flow*. Similarly, if *ϑ* = 0*^◦^*, the flow is assisting the movement of the individuals towards the target and is described as *favourable flow*. We use the term *cross flow* for the case *ϑ* = 90*^◦^* or 270*^◦^*, and the remaining cases are termed *offset flows*.

For the real-world flow we set:

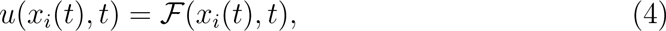

where *F* denotes a realistic, turbulent, time-dependent flow field. Specifically, this flow field is obtained from HYCOM data, (hycom.org, see Supplementary Information D). The flow velocities are subsequently normalised such that the mean flow strength (averaged over space and time) equates to the fixed flow strength parameter *ζ >* 0.

All simulations and figures were produced in Julia and the code is available online at https://github.com/Tom271/CollectiveNavigation. Table 2 summarises the model parameters and notation used.

**Table 2:**
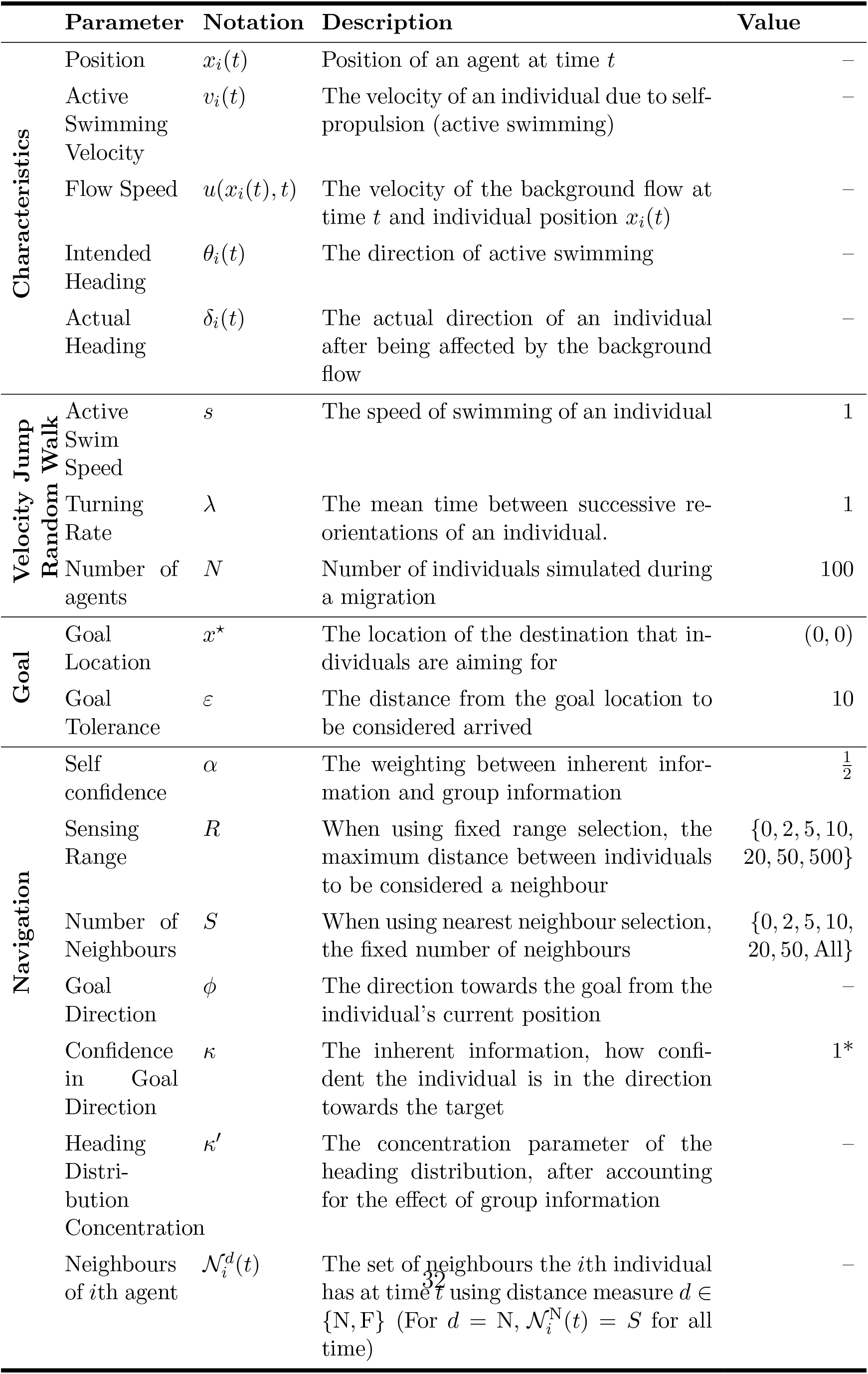
Notation used and parameters of the model. An en dash (–) indicates a value that potentially varies in space and/or time. (*) See Supplementary Information C for simulations under variations to these values.

## 3 Results

This section is structured as follows. We first ensure that our model recapitulates well-known results about: i) the effectiveness of collective navigation over individual navigation in a non-flowing environment (e.g., [31]); ii) the impact of flow on the ability of a solo navigator to reach some target destination (e.g., [33]). Following this, we explore how the different forms of flow impact on collective navigation for the various submodels summarised in Table 1. Animations of select simulations are available online at https://tom271.github.io/Supplementary-Information/.

### 3.1 Base Scenarios

#### Collective navigation is beneficial, but only above a critical level of group information

In Figure 2a) we present the mean (*±* one standard deviation) of the median arrival time obtained from 50 realisations of the [FI] model in the absence of flow. Here, intended and actual directions coincide and, consequently, the results will be equivalent for the [FA] model. We note that the median time of arrival first *increases* as the sensing range increases from *R* = 0 (no collective navigation) until *R* ≳ 5.0. The median arrival time then decreases, where beyond *R ≈* 20.0 we see little improvement in efficiency of navigation (see also Figure S1). This is analogous to the results seen in [10, 31], where groups are observed to arrive nearly 20% faster for large levels of collective navigation than in the individual case. For intermediate sensing ranges, the variance in the arrival time is increased when compared with the extremes of sensing range (see Figure S1). This is due to there being more possible arrangements of subgroups of few individuals than when the whole group is interacting, or when individuals navigate alone. Overshooting the goal in these regimes can also result in a delay due to information propagation through the group. When an individual overshoots the goal, the next time it reorients it is most likely to head back towards the goal. In a collective navigation scenario, however, others in the group may still be swimming away from the target. When a group member reorients in this setup, it continues to be influenced by the overall group moving in the wrong direction. This is more prevalent in a larger group as it takes longer to influence the group’s direction. This is mitigated, though, by the fact that a large group is less likely to overshoot due to improved navigation. Note that for the study here, where the initial configuration considers a population distributed uniformly across a square with side length 40, there is minimal difference in the dynamics beyond *R* = 50. At higher sensing ranges, although the bulk of the population arrives quickly, we note there is a longer tail to the arrival time distribution. Effectively, as more of the population arrives, remaining navigators lose the benefits of collective navigation. In the [NI] and [NA] models, very similar behaviour is seen in the absence of flow, with a slightly more noticeable reduction in performance at a lower number of neighbours.

**Figure 2:**
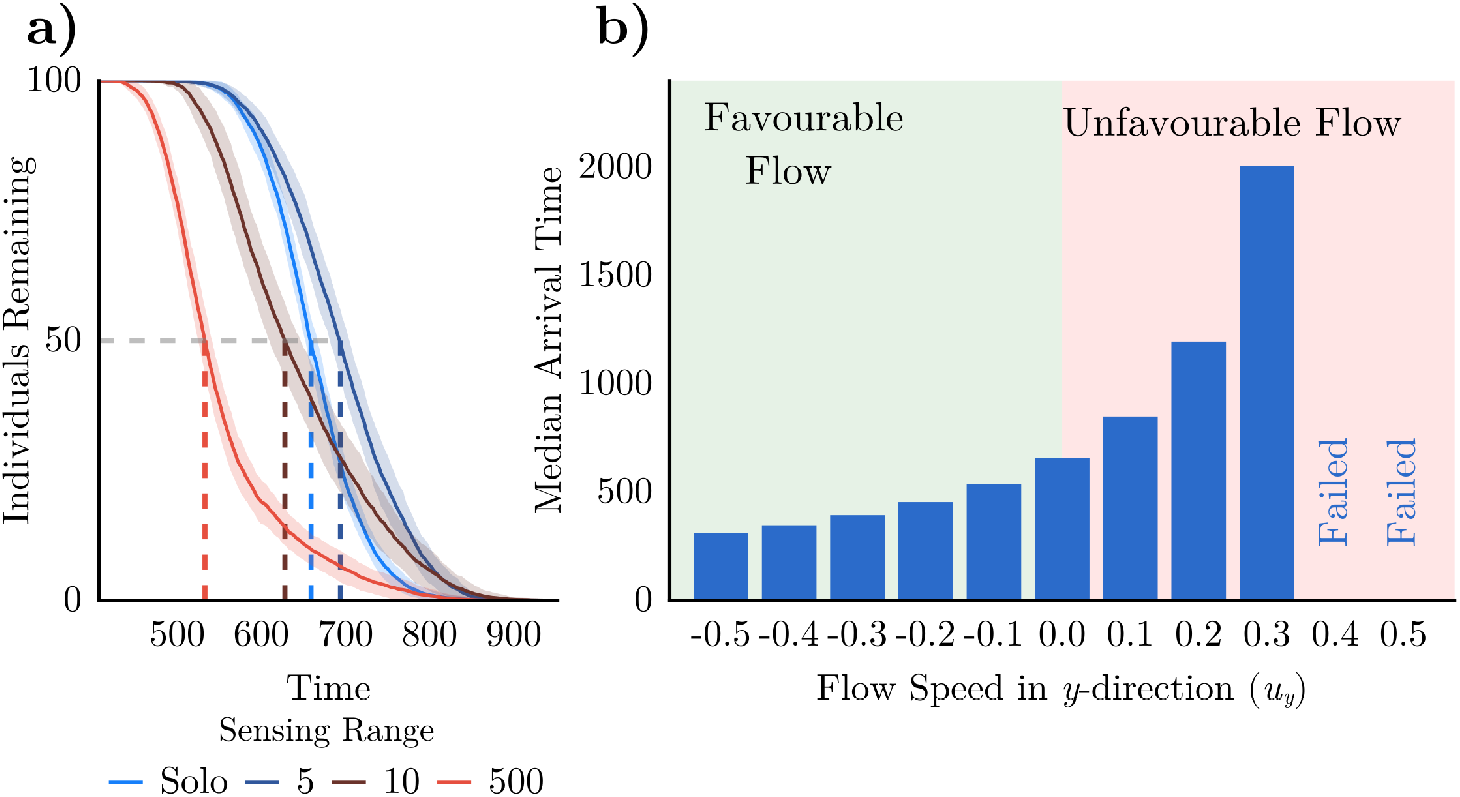
Collective navigation is beneficial above a certain information threshold; opposing flows impede success for an individual. a) The number of individuals yet to arrive for different sensing ranges for the ([FI] /[FA]) models in the absence of flow (*ζ* = 0). Solid lines show the mean, surrounding ribbons show the standard deviation from 50 realisations. Dashed lines show the median arrival time. Sensing ranges beyond *R* = 20 show similar behaviour to *R* = 500. b) The median arrival time averaged over 50 realisations, as a function of flow speed in the *y*-direction (*u_y_* = *ζ* sin(*ϑ−* 90*^◦^*)), for a population of individual navigators. Beyond a critical flow strength, migration fails to occur i.e., less than 50 out of 100 individuals arrived within the simulation time.

#### Flow can facilitate, inhibit, or even prevent individual navigation success

In Figure 2b) we show the impact of a simple flow (Equation 3) on a population of non-communicating individuals. A flow parallel to the migration direction is varied in strength from *ζ* = 0 to 0.5 in increments of 0.1 for *ϑ* = 0*^◦^* and *ϑ* = 180*^◦^*. Recall *ϑ* = 0*^◦^* indicates flow in the direction towards the goal and hence a favourable flow. Unsurprisingly, for the favourable flow, we see faster arrival times. Conversely, for an unfavourable flow, *ϑ* = 180*^◦^*, the migration time increases with *ζ*, until the point of failure when *ζ* ≳ 0.3 (a flow speed 30% of the individuals’ active movement speed). Intuitively, higher flow strengths limit the effective swim speed of an individual – for a more detailed study, see [34], where the flow speed at which failed migration occurs is determined according to the certainty in the target direction. Note that the behaviour here will be the same for all variants of the model, as no group information is included.

### 3.2 Simple Flows

#### Above a critical sensing range, sharing intended headings is universally beneficial under simple flows

In the [FI] model, under non-zero flows, group-level behaviour depends on the flow strength. For low sensing ranges, we observe behaviour very similar to individual navigation (Figure 3a). Here, the sensing range between individuals is insufficient to maintain contact and members effectively navigate in isolation. When the sensing range is sufficiently high (*R* ≳ 20), individuals can maintain contact with others throughout migration, and hence continue to receive information that reinforces navigation. In a favourable flow, the median arrival time reduces irrespective of the sensing range. This is expected since individuals will have a higher effective swimming speed due to the agreement between the active swimming velocity *v_i_*(*t*) and the flow velocity *u*(*x_i_*(*t*)*, t*). Across all studied flow strengths, the effect of increasing the sensing range mirrors that observed in the absence of flow case.

**Figure 3:**
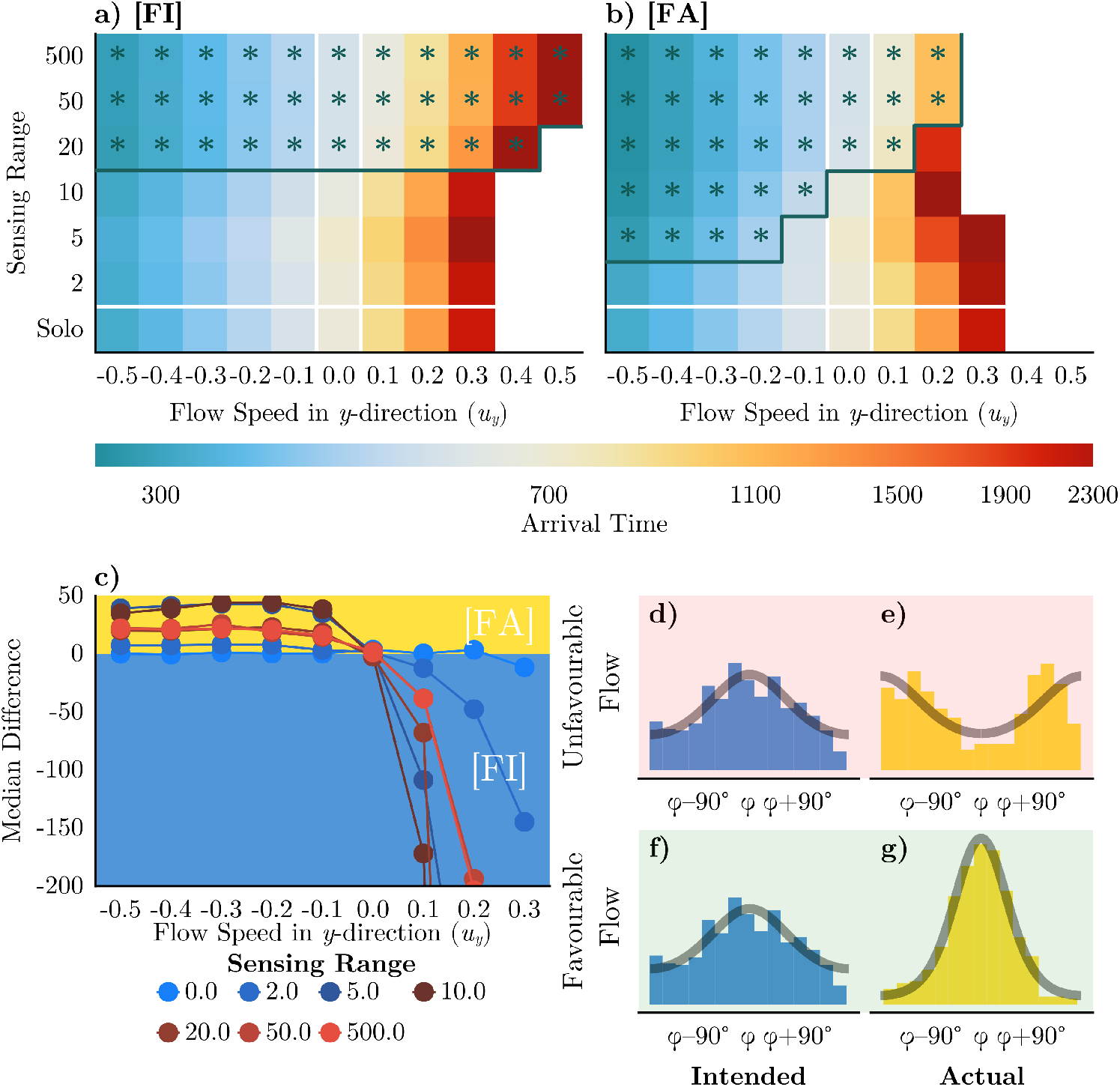
Effect of Perception Type depends on Flow Direction. (a-b) Heatmaps of the median arrival time as a function of the flow speed in the *y*-direction and sensing range. Each parameter combination is averaged over 50 realisations. a) Results for a fixed range interaction and intended headings [FI]. b) fixed range interaction and actual headings [FA]. Green asterisks denote parameters where solo navigation was outperformed by *≥* 5%. The green solid line indicates the boundary of the region where collective navigation induces a *≥* 5% improvement. Missing cells indicate where there is failed migration. c) The difference in median arrival time between the [FI] and [FA] model across a variety of flow strengths and sensing ranges. Positive values (yellow) indicate scenarios in which sharing actual headings leads to faster arrival times than when sharing intended headings (blue). (d)-(g) Illustrative neighbour heading distributions relative to the goal direction *ϕ* in unfavourable (d-e) and favourable (f-g) flows.

#### Collective navigation can allow a group to arrive in a scenario where individuals generally fail

The benefits of collective navigation are particularly prominent when swimming in an unfavourable flow regime, where we observed that individual navigation fails above a critical flow strength. While low sensing ranges (e.g., *R* = 2) yield equivalent behaviour to zero communication, and moderate sensing ranges (e.g., *R* = 5) lead to even poorer performance, larger sensing ranges (*R* ≳ 20) can provide a level of benefit that permits successful navigation in cases where individual navigation fails. Failed migration is the inevitable consequence of very strong unfavourable flows. However, for group navigation the critical flow strength for failure is extended and a group can overcome stronger flows.

In unfavourable flows, the first group failures are observed. As the sensing range increases, performance improves – at sufficiently high collective information and unfavourable flow (*ζ ≈* 0.1), a group can even beat the performance of non-communicating individuals in a static environment. We note that the arrival time distributions have much higher variance, due to the lower effective swimming speed of the group (see Figure S2 in the Supplementary Information). When swimming against the flow, few individuals reach the target unless the sensing range is sufficiently high. The benefits of collective navigation are therefore particularly pronounced when swimming in unfavourable flows.

#### In favourable flows, sharing actual headings is more effective than sharing intended headings

We consider a performance comparison according to whether an individual receives intended or actual headings from neighbours. In Figure 3b) we summarise the data from simulations of the [FA] model. While the [FI] and [FA] models are equivalent in a stationary environment, they yield subtly distinct behaviour in the presence of flow. For a favourable flow, perceiving the actual headings is more beneficial than perceiving the intended heading. This follows from the fact that actual headings are more aligned with the goal direction than intended headings. In the [FA] model, the flow has a positive effect irrespective of sensing range. In Figure 3f-g) we illustrate this, as we see that actual headings are more concentrated in the target direction and are thus an aid to navigation. Larger sensing ranges allow for the collection of a greater number of concordant headings, and thus create a larger improvement over the intended heading case, see Figure 3c).

#### In unfavourable flows, sharing actual headings is highly detrimental

In stark contrast to the advantages observed for favourable flows, for unfavourable flows, perceiving the actual headings is highly disadvantageous. Here, actual headings tend to be less aligned with the goal direction than intended headings, and hence there is less certainty in the received collective information. Note that this effect can become extreme to the point that any level of collective information may be detrimental, and it becomes more advantageous to navigate as a solo navigator. In Figure 3e) we see that actual headings generally point away from the goal and any level of collective information acts against the individual’s inherent knowledge of the target direction. In Figure 3c) we summarise these differences between the [FI] and [FA] models and show that the [FA] model is near-ubiquitously advantageous for the favourable flow, but near-ubiquitously disadvantageous for an unfavourable flow. In Supplementary Information Figure S5 we show the same heatmaps, but annotated with median arrival times relative to the solo navigator case, for both intended and actual headings.

#### For cross and offset flows, sharing intended headings is the more robust strategy

We extend our investigation by considering a range of cross and offset-flows. First, we consider cross-flows of increasing strength (Figure 4a-c). In the absence of flow the two models are equivalent and trajectories are reasonably direct to the goal (Figure 4a). A moderate cross-flow, however, generates a detour which becomes significantly longer under shared actual headings (Figure 4b). Stronger flows lead to an even more marked contrast, where perceiving intended headings can still allow successful goal navigation, but perceiving actual headings leads to failure (Figure 4c). Here, the group becomes lost by sharing effectively low quality information that subsequently reinforces the wrong direction due to its incorrect mean.

**Figure 4:**
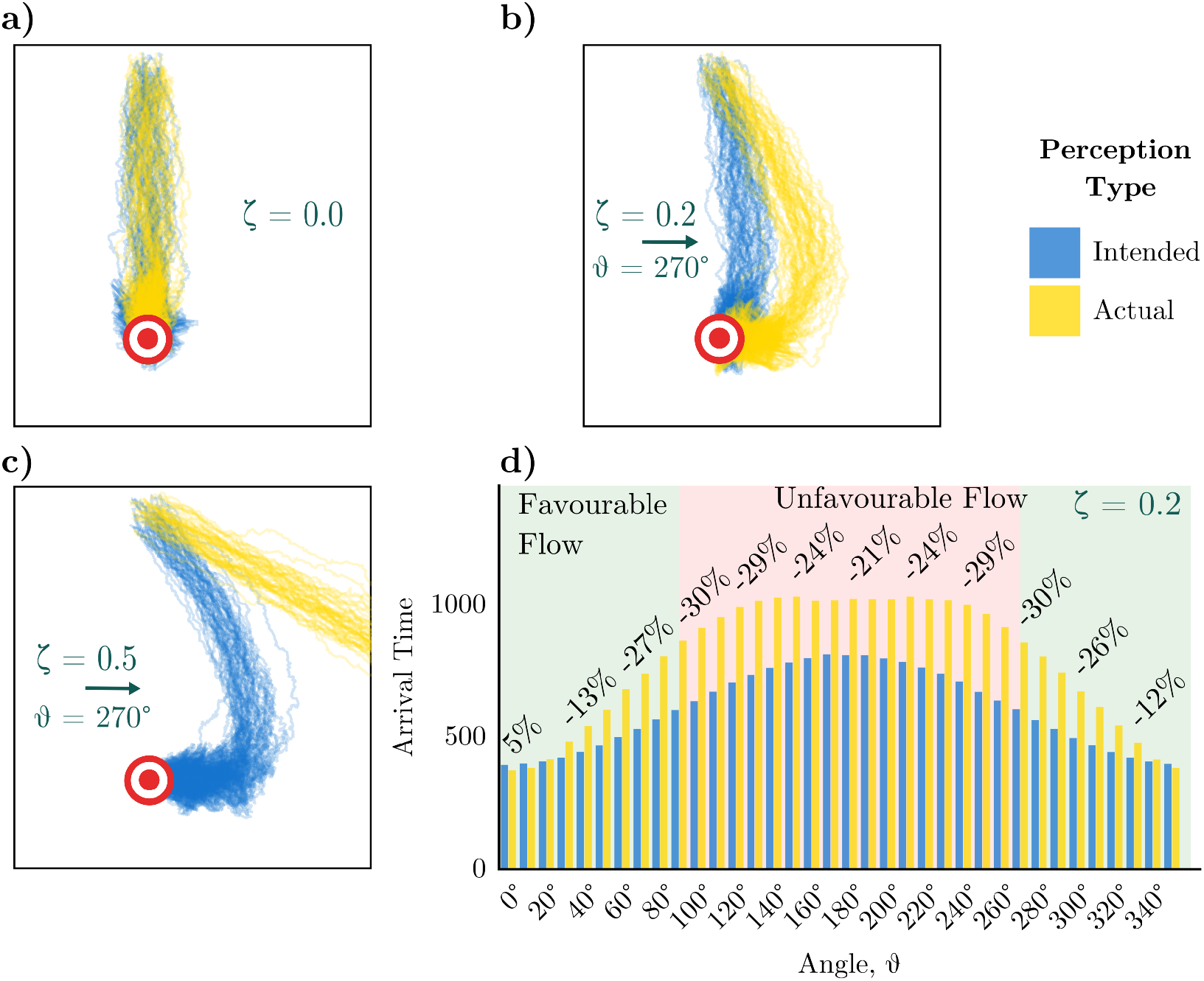
Perceiving actual headings can mislead the group resulting in slower arrival times across many flow angles. (a-c) Trajectories of 100 individuals navigating in cross flows of varying strengths in the [FI] (blue) and [FA] (yellow) models. The goal is represented by yellow and green concentric circles. d) The effect of varying the flow angle *ϑ* with fixed flow strength (*ζ* = 0.2) on the median arrival time for the [FI] (blue) and [FA] (yellow) models, averaged over 50 realisations. Annotations show the percentage difference in median arrival time.

We next consider offset flows (Equation (3)), i.e., where the flow angle varies with respect to the axis of straight-line navigation. Specifically, we consider a fixed sensing range *R* = 20 and fixed strength of flow (*ζ* = 0.2), but vary the angle of flow in increments of 10*^◦^* from 0*^◦^* to 360*^◦^*. Angles close to 0*^◦^* (or 360*^◦^*) represent broadly favourable flows, while angles close to 180*^◦^* represent broadly unfavourable flows. As indicated in Figure 4d), a relatively narrow window of angles exists under which perceiving the actual headings is advantageous, with up to 5% improvement in median arrival times observed, relative to perceiving intended headings. Contrast this with the worst case scenario, where the arrival time can be up to 30% slower when perceiving the actual headings. This reinforces the notion that perceiving intended headings is broadly more advantageous.

#### Navigation is robust with respect to the method of neighbour selection

Figure 5 summarises the same sequence of simulations as in Figure 3 performed when individuals interact according to the [NA] and [NI] models. Results are broadly in line with those of Figure 3, suggesting that the type of communicated information (actual or intended) is more influential than the selection of neighbours (fixed range or nearest) for navigation success. Note that there is no direct correspondence between the number of neighbours (*S*) and the fixed sensing radius (*R*): for the fixed sensing radius models the number of members of the interacting group changes throughout navigation, whereas for nearest neighbour models it remains constant (until there are less than *S* individuals remaining). Despite this, generally, we observe similar performance between nearest neighbour and ranged interactions: worse performance for a low number of neighbours, and improved performance at higher numbers. Further, actual headings are slightly advantageous for favourable flows, but highly disadvantageous in unfavourable flows. Note that when interacting with a fixed number of neighbours, the amount of information is capped; which can be disadvantageous when in a dense group as information is lost from nearby individuals. In an unfavourable flow with low strength, we see slightly worse performance when navigating using a fixed number of neighbours. In an unfavourable flow, individuals are pushed apart due to the increased variance in the heading distribution caused by the flow. In a favourable environment, this does not occur: the heading distribution is usually more tightly concentrated around the goal direction. This initial dispersal means that under the ranged interaction, when the sensing range is low, the initial group promptly spreads and individuals fall out of sensing range. When using nearest neighbour communication, the individuals retain communication with their neighbours, irrespective of dispersal. Counter-intuitively, this results in worse navigation as the information they receive from neighbours is often dissonant.

**Figure 5:**
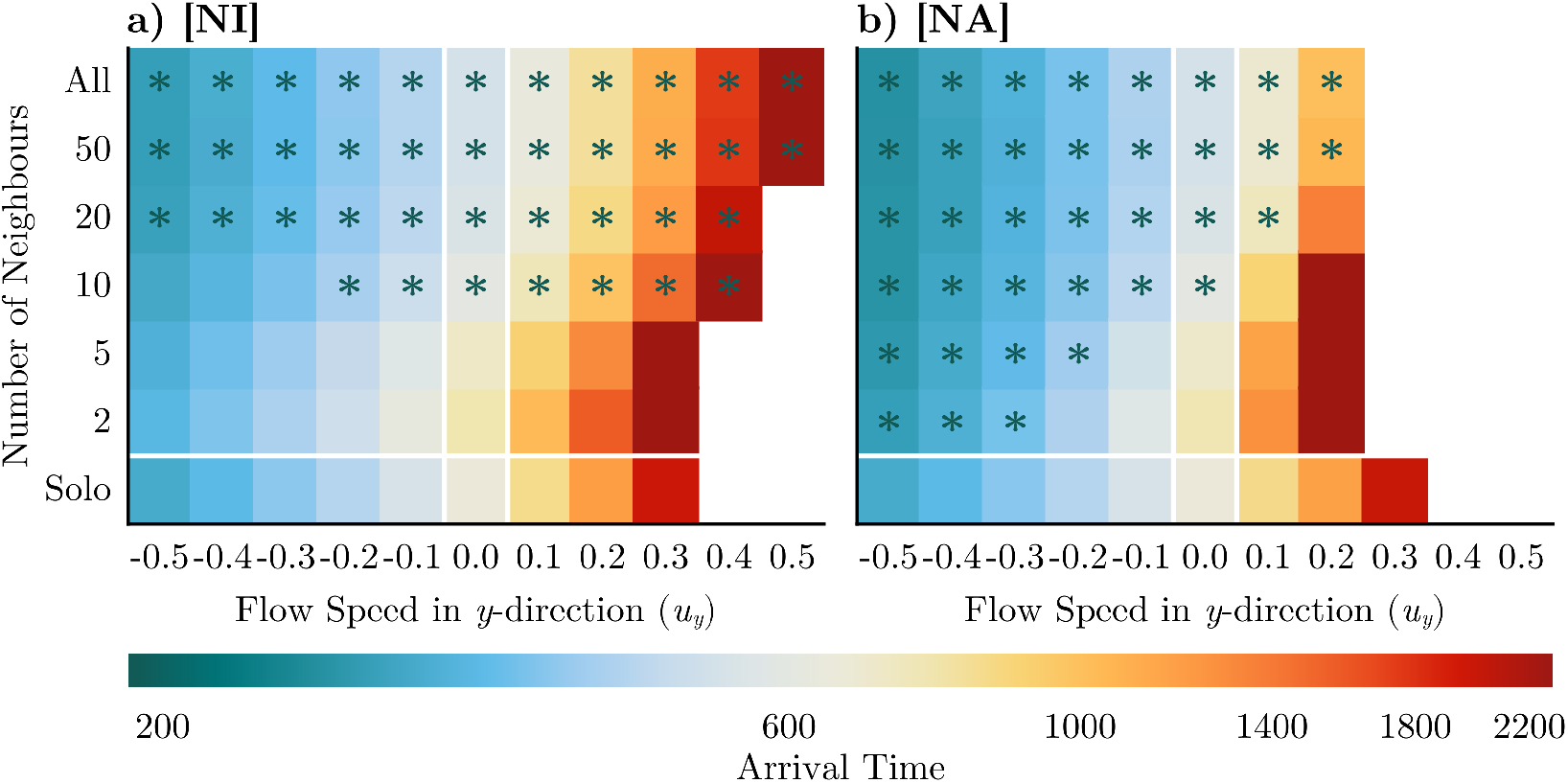
Neighbour selection method has little impact on effect of collective navigation. (a-b) Heatmaps of the median arrival time as a function of sensing range as in Figure 3. a) [NI] model. b) [NA] model. All other parameters remain unchanged. Green asterisks denote regions where the migration was more than 5% quicker than in the solo navigator case, while the solid green line marks the boundary of this region.

The discrepancy in performance is due to the average group size. When an individual navigates in the absence of others, the concentration parameter of the heading distribution is fixed at *κ^′^* = 1. If instead the group is collectively navigating, this concentration parameter can increase or decrease (as can the accuracy of the location parameter). If the group is heading towards the target (which is more likely in a favourable flow), information is beneficial and thus the highest group size is rewarded. In contrast, when navigating against the flow, group information is a disadvantage as it likely more dissonant, and thus a lower group size is better.

### 3.3 Real-World Flows

The previous section provides evidence for the benefit of collective navigation in linear flows, but such regular flows are rare in natural environments. In this section, we apply a similar analysis to the real-world flow field described in Section 2.3.

#### In real-world flows, sharing actual headings is detrimental

Above, we observed that for almost all angles within an offset flow setting, collective information based on intended headings outperformed that based on actual headings. Real world flows will rarely align with the goal direction and thus we observe a similar detrimental effect when collective information is based on actual headings. In Figure 6a-d) we show heatmaps for the median arrival time under the four model forms. We see that [FI] and [NI] formulations outperform their [FA] and [NA] counterparts under all flow strength and interaction range combinations. Moreover, [FA] and [NA] models reveal a drastically reduced region in which successful navigation can occur. Under intended headings, collective navigation can be beneficial across a reasonable spread of flow strengths and communication ranges. However, navigation based on actual headings is detrimental to the point that it is often advantageous, in terms of travel time, to be a solo navigator. Here, an increased sensing range is universally unhelpful due to the more varied information provided by actual headings under disordered flows. To provide a more detailed picture for lower flow regimes, we also provide the explicit data for the [FI] and [FA] models for smaller increments of flow in Figures 6e-f). Here we observe that sharing actual headings can still provide some benefit to navigation, but only if the flow is relatively weak compared to the active movement speed and the sensing range is sufficiently high. However, the benefit of sharing intended headings is much higher, and the region larger. Beyond a moderate flow strength, navigation failure occurs for all migration attempts that utilise group information. At this point each individual is heading in quite different directions, and each reorienting individual is flooded with less relevant information. As previously, we note that navigation is largely robust to the neighbour selection method. In Figure 6a-d) we see, again, the type of information that is communicated has more impact than the neighbour selection method. Figure S6 shows heatmaps annotated with median arrival time relative to the solo navigator case, for all submodels.

**Figure 6:**
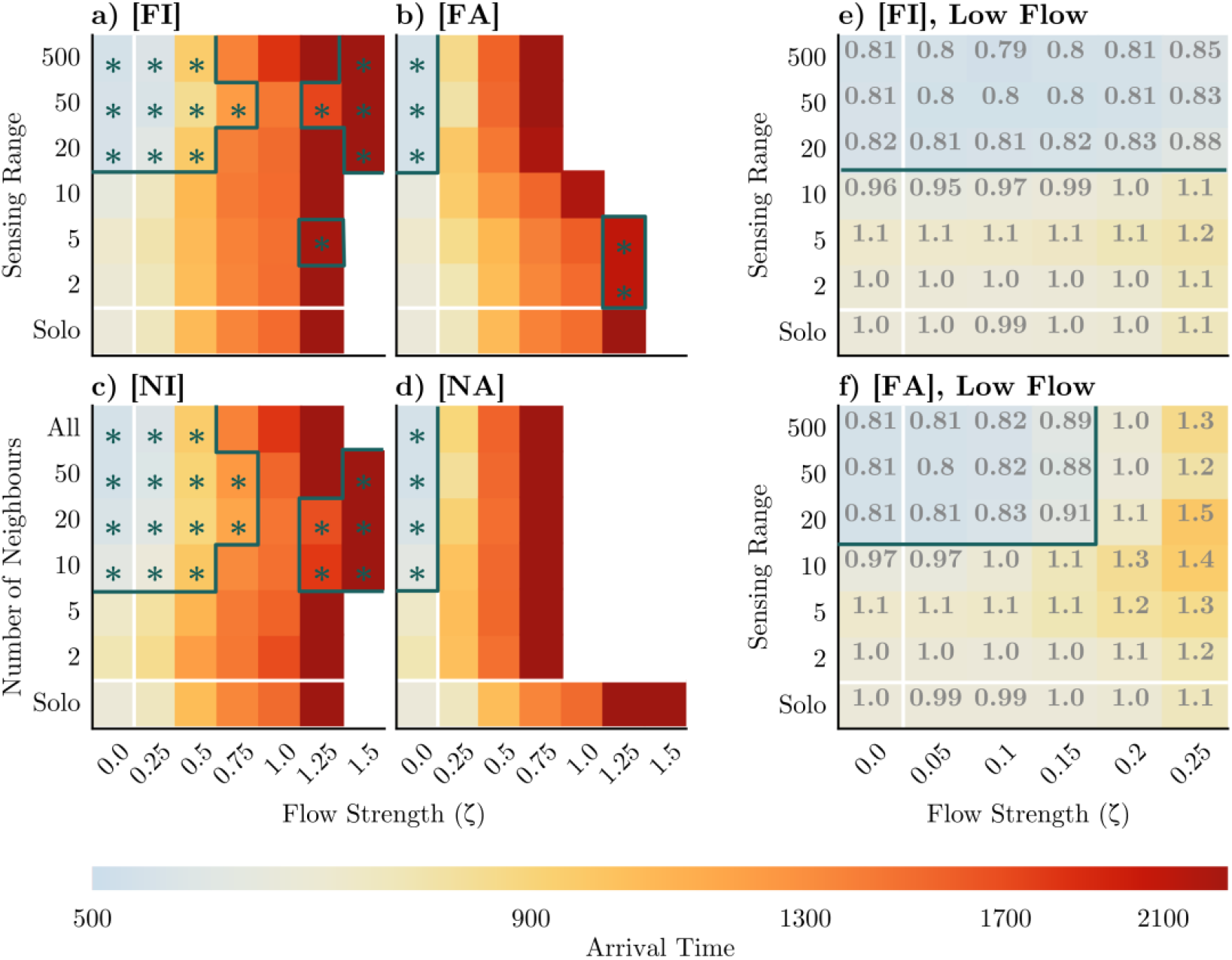
Perceiving actual heading is detrimental to migration success in a turbulent flow. (a-d) Heatmaps of the median arrival time relative to an individual in the absence of flow, averaged over 25 realisations, for the [FI], [FA], [NI] and [NA] models respectively. All simulations are in a turbulent flow environment. The grey dashed line indicates the region expanded in Figures (e-f). (e-f) The median arrival time relative to the solo navigator case for a range of lower flow strengths. Grey annotations indicate the time taken relative to a solo navigator in flow. The colorbar is shared across all axes and Figure 5.

#### There exists an optimum level of group information for success in a turbulent flow

In Figure 6a) we show a broadly similar pattern to those observed for the laminar flows (cf. Figure 3a)): specifically, higher sensing ranges improve efficiency of navigation. However, in contrast to the laminar flow case we observe that at the highest sensing ranges the navigation becomes slower: a phenomenon that persists across all flow strengths tested. When individuals become separated by a large distance, their experience of the flow and the goal direction are likely to be very different. In this scenario, group information is unhelpful. Therefore, while collective navigation is beneficial if individuals communicate intended headings, this only remains the case if the individuals communicate across a particular range: large enough to obtain a sufficient quantity of good information, but not so far as to start acquiring irrelevant information due to the turbulent nature of real-world flows.

## 4 Discussion

Numerous animal species perform migrations as a group, often when immersed in a complex flow. In such an environment, each individual’s actual heading may differ from its intended heading, due to passive drift. Accordingly, it is unclear as to how any collective navigation benefits are altered according to the perceived heading. We have directly addressed this by extending a previous model of collective navigation [31] to include flow and different mechanisms of group sensing/information transfer. Under simple unidirectional flows (e.g., river environments) the better strategy depends on the flow direction: when navigation is strongly with the flow (e.g., down a stream) it is more beneficial to receive actual headings; for nearly all other flows, the intended heading is more beneficial. Under the convoluted, turbulent, and strong flows often encountered in nature, collective navigation is only beneficial when an intended heading is perceived. Navigating according to actual headings is often worse than when performing a solo navigation. These results are robust to the choice of neighbour selection method. That is, whether neighbours are defined as the population up to some fixed distance (as in [11, 26]) or up to a fixed number of nearest neighbours regardless of their distance (as in [21]). Our results align with previously observed breakdowns of the ‘many-wrongs’ principle such as in [10], where failure was induced by adding noise to the orientation of individuals. We have shown here that any benefits of group navigation may break down when navigating in a turbulent flow, according to the communicated information.

Whether actual or intended headings are perceived will, critically, vary according to the species, context, and environment. Intuitively, collective navigation based on actual headings is the more simplistic assumption: no ‘willing’ communication is required by the navigators, as other individuals could obtain this through direct observation (e.g., visually tracking the path of the neighbour). Intended headings, on the other hand, could suggest some higher-level communication if conspecifics are out of sight: e.g., a neighbour audibly communicating its intended path. We studied the two extreme cases: communicating only intended or only actual headings. In reality, it is possible that the information may lie somewhere between these two extremes. While human navigators can communicate complicated information, the degree to which this is possible in animals is largely unknown. Species of whales, renowned for the complexity of their calls and song [20], may offer an example of a species capable of communicating detailed information. The context of the migration is also a factor to consider. In a cooperative scenario, an individual may freely communicate their intended direction to aid navigation success; whereas, in a competitive scenario, individuals may suppress active broadcasting of their intended route. Our results show that in instances where only actual headings are or can be perceived, it is often disadvantageous to utilise collective navigation when it comes to minimising navigation time. Group structures may still of course have other benefits, e.g., reducing the threat of predation [35].

The extent to which collective information improves or hinders navigation fundamentally depends on whether confidence in the target direction is increased or decreased. Actual headings only improve navigation when flow is aligned with the goal, as it is only within that (restrictive) scenario in which headings become more concentrated towards the goal. Intended headings offer more robust benefits, even for highly turbulent and strong flows: here, individuals always relay their knowledge of the goal according to inherent information, which is unaffected by the flow. The quantity of information is generally beneficial, in that obtaining the headings of more neighbours is generally beneficial, as there is an increased reduction in the uncertainty. However, a note of caution is required here as, under variable flows, unrestricted gathering of information becomes detrimental. Once an individual is too distant, it has a highly distinct intended heading, and the relevance of its associated information is reduced. In other words, while quantity is important, quality of information trumps quantity.

The quality over quantity issue highlights an ingredient missing from our current model that merits closer inspection. Specifically, we have no explicit aggregating effect within the model, as orientation is based purely on the combination of inherent information and alignment of headings. This does help maintain aggregation – as individuals tend to migrate in a common direction towards a target – but it does not explicitly stop individuals moving out of communication range. Adding an aggregating mechanism, e.g., through separate zones of attraction/alignment [11, 27], may mitigate against excessive dispersal by reorienting wayward individuals back to the fold. It should be noted that whether this helps navigation is not clear cut: any centralising tendency may reduce the time spent swimming towards the goal, increasing overall navigation time. Thus, examining the potential tradeoff between maintaining a compact group structure that ensures individuals have common target directions, and solely aiming for the goal merits further attention.

A key question studied here is based on the observation that, in flowing environments, intended and actual headings are generally unaligned due to passive drift. A potentially significant factor that has not been addressed here is the extent to which flow-alignment and flow-compensation mechanisms factor into a movement response. Many aquatic and airborne organisms have a capacity to detect flow and align their body axis accordingly: an orientation response known as rheotaxis (for water currents) [36] or anemotaxis (for air currents) [14, 15, 16, 37]. Assuming a capacity to actively or passively detect a local flow may allow an individual to partly compensate for excessive drift, narrowing the difference between intended and actual headings. In our model, such responses could be brought into the modelling framework through an additional component within new heading selections that depends on the external flow vector. However, incorporating compensation is non-trivial. If we assume an individual factors flow into its decision making, the chosen active velocity will combine information about the flow and the goal, resulting in a further communication type to be considered (see Figure 7). Under a variable flow further subtleties will arise, such as the temporal and spatial scales over which an individual could detect the flow angle and strength: can an individual calculate some ‘average flow’, or does it only have instantaneous knowledge? A full study of the (possibly positive or negative) impacts of flow compensation on group navigation is certainly warranted, and is left for a future work dedicated to this issue.

**Figure 7:**
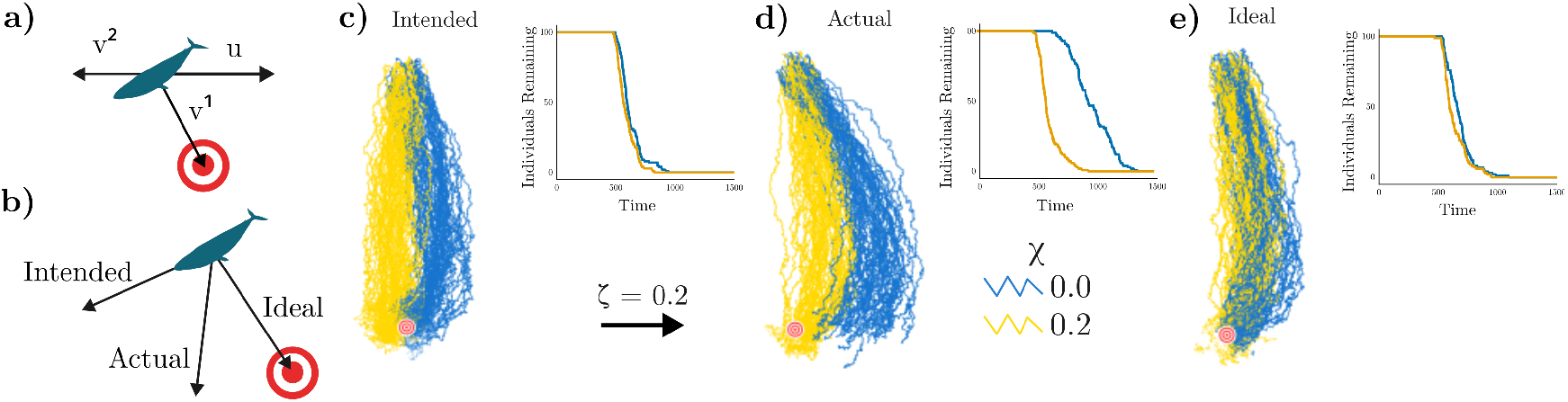
Complications of Drift Compensation. One method to incorporate drift compensation would be to split the active swimming velocity (*v*) into two components *v*^1^ and *v*^2^, where *v*^1^ is the active swimming velocity dedicated to swimming in the goal direction (*ϕ*), and *v*^2^ is the portion of active swimming velocity dedicated to compensating for the flow, see a). The model becomes 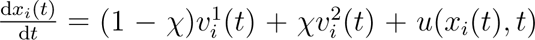 where 0 *≤ χ ≤* 1 is a parameter controlling the amount of effort dedicated to compensating for the flow rather than seeking the goal. Consequently, this introduces further subtlety to communication, as it is now possible to communicate: *ideal* direction (arg (*v*^1^)), intended direction (arg (*v*^1^ + *v*^2^)) or actual direction (arg (*v*^1^ + *v*^2^ + *u*)), see b). c-e) show the impact of these different communication types for a simple cross flow with *ζ* = 0.2*, ϑ* = 90*^◦^*. Blue trajectories are those for a navigating population with no drift compensation (*χ* = 0), while yellow trajectories are those under a slight compensation (*χ* = 0.2). All inset plots show the number of individuals yet to arrive at the goal against time.

Our current work has intentionally focussed on an abstract setting, i.e., exploring the general benefits and costs of collective navigation in a flow for an unspecified animal species. As such, the model was formulated in a non-dimensional setting. Specifically, we considered speed and movements rescaled to arbitrary length units and a distance to target migration of *∼* 300; effectively, a macroscopic setting in which each individual must sample their surroundings (on average) hundreds of times before reaching the goal. The real-world flow data was also recast onto the non-dimensional space and time scales, parameterised by a single ‘flow strength’ parameter that represents the average flow speed in the region. A value *ζ* = 0.25 indicates the average flow is 25% of the individual’s mean active movement speed. To provide further context to these numbers, it is helpful to consider the strengths of real-world flows relative to actual movement speeds. For example, for the flow data considered in Section 3.3 the dimensional mean flow speed for this region is 0.94 km h*^−^*^1^, and hence a value of *ζ* = 0.25 would correspond to the flow experienced by an animal with sustained movement speeds of 3.76 km h*^−^*^1^. At these values, it suggests that collective navigation based on the intended heading could provide up to 25% improvement over solo navigation. In contrast, utilising information based on actual headings in this regime provides a 30% *decrease* in performance, see Figure S6. To take one example, the Atlantic salmon, *Salmo salar*, has an optimal speed (in the sense of minimising energetic cost of transport) of around 2.5 km h*^−^*^1^ [38]. In our model, this corresponds to *ζ ≈* 0.4, a region in which sharing the intended heading gives clear advantage over sharing the actual heading (Figure 6a-b). In contrast, blue whales, *Balaenoptera musculus*, can maintain speeds of *∼* 6 km h*^−^*^1^ during migrations [39]. This corresponds to *ζ ≈* 0.15, a region in which the benefit of collective navigation is clear regardless of perception type (Figure 6e-f).

In this study we have focused on the median arrival time as a measure of migration success. This is chosen for its robustness and insensitivity to the long tails of the arrival time distributions, befitting of a study conceived in a general setting. For a more specific modelling study, the appropriate metric could significantly differ, as alluded to in Section 2. For example, when bird species migrate to a nesting site, the first to arrive will be able to choose the best nest with the quality decreasing until no suitable sites remain [40]. Consequently, the time of first arrival may be a more suitable metric. For species that form strongly social and familial structures or have few offspring, a driving aim would be for the majority or all the migrating group to arrive [41]. Within this context, the behaviour of ‘successful’ navigators also becomes important. Here we have assumed individuals that have arrived at the goal no longer play a role, perhaps befitting of ‘selfish social navigators’ that do not particularly care about the population once they have arrived. Another interesting assumption would be for arriving navigators to strongly signal their success, which intuitively would help round up stragglers and avoid long tails in the arrival time distribution. In many cases, the goal location may not be fixed in position, such as a food source actively moving or drifting on ocean currents [42, 43]. An alternative approach that avoids this complication is that of [28, 8], in which the goal is placed ‘at infinity’ and the navigational efficiency is calculated rather than arrival time. This has been considered in Supplementary Information A, where we see that utilising actual headings confers even greater benefits under favourable flows. Navigation using intended headings, however, remains the more robust strategy.

According to the environment and method of communication, signal propagation may also be impacted by the background flow. As an example, when communicating via sound in an ocean, the signal propagation is less affected by turbulence due to the speed of sound propagation in water, however surface and seabed effects become relevant [44]. In aerial migrations, however, sound may be heavily affected. Incorporating a position-dependent noise term into the model could account for this. This could also also account for a more general noise in the information environment, which has remained fixed in this study.

Beyond animals, navigation is also of clear interest in robotics, to develop autonomous remote-controlled agents able to navigate in complex flowing environments [45]. This is seen in subsea remotely operated vehicles (ROVs) in the extractives industry, as well as in drone flights. The question of route-finding in flow is an obvious optimisation problem and can find its (mathematical) roots in Zermelo’s Navigation Problem [46]. Zermelo himself solved the question in the constant flow case and showed that the general case is the solution of Zermelo’s equation, a nonlinear PDE derived from the Euler-Lagrange equations. Solving this problem in a turbulent setting has attracted recent interest using the tools of reinforcement learning, see [47, 48].

Animal groups are, of course, typically far from homogeneous. Heterogeneity could enter the group in numerous ways: genetic variation affecting, say, swimming or flight speed, [49]; pre-migration history leading to navigators with differing fitness or energy reserves [50]; social structures within groups, e.g., based on age, maturity or experience [51, 52]. Incorporating such heterogeneity, of course, would require careful consideration: for example, more mature individuals may have access to higher levels of inherent information, and could also have a greater weighting in the collective information process of others; fitter individuals could have differing swim speeds, and so on. Indeed, a strength of the modelling framework is that it allows incorporation of heterogeneity in a myriad of ways. Accordingly, the framework could be potentially targeted to address highly specific instances of migration.

Another source of navigation information that has not been considered here is *inertial information*. Individuals may use their own movement history to inform future movement, as accounted for in correlated random walk models, e.g. [32, 8]. This history could also be included for conspecifics, whereby an individual remembers the positions of neighbours to determine their headings.

Throughout this study, we have assumed that the self-confidence parameter, *α*, was fixed throughout the population, and in time and in space; indeed, relying on previous sensitivity analyses, this was set at a ‘broadly optimal’ value [31]. More generally, we would expect self-confidence to be highly dynamic, e.g. selected according to the level of inherent information (*κ*) or commonality of co-navigators alignment: it is reasonable to assume that individuals have periods in which they are confident in their inherent information and see no value in group information, while in patchy information environments there may be benefit in aggregating and trusting a group more [31]. Extending the model to include dynamic self-confidence would form a natural extension.

Advances in remote sensing technology are bringing novel data sources to the study of animal migration, allowing tracking of species within highly complex and inaccessible environments such as the ocean at ground-breaking spatiotemporal resolutions [53]. Utilising this data, coupled with ocean forecasting models, is providing new insights into how animals behave on long distance migrations. The study here has shown that while näıve collective migration based on actual headings could be counterproductive in terms of median migration times, the use of intended headings can still confer significant advantages. Bridging the gap between the abstract modelling approach taken here and specific animal migrations through the use of high-resolution data could provide insights into the anthropogenic impact on migrations, allowing for science-informed policy that balances human activity and animal welfare.

## Data Access

The code used to implement the models is available at https://github.com/Tom271/CollectiveNavigation. The lookup table is available at https://melbourne.figshare.com/articles/dataset/kappaCDFLookupTable_mat/14551614.

## Authors’ Contributions

T.M.H. conceived the study, developed the modelling framework, designed and performed the numerical experiments, designed and performed the analysis, and wrote and edited the manuscript. S.T.J. conceived the study, developed the modelling framework, designed the numerical experiments and edited the manuscript. K.J.P. conceived the study, developed the modelling framework, designed the numerical experiments, designed and performed the analysis, and wrote and edited the manuscript. M.O. developed the modelling framework, designed the numerical experiments, and edited the manuscript. All authors gave final approval for publication.

## Acknowledgements

This work has made use of the resources provided by the Edinburgh Compute and Data Facility (ECDF) http://www.ecdf.ed.ac.uk. We would like to thank the reviewers for their helpful comments that have greatly improved the manuscript.

## Footnotes

1 Possibly the only non-human to have a multi-platinum record [19]

2 The modified Bessel function of the first kind of order zero can be defined as 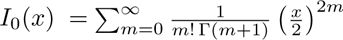 where Γ is the Gamma function.

3 In some literature, the intended heading is known simply as the *heading* while the actual heading is the *track* [13, 16, 14]

