## Supplementary Information for "Intent matters: how flow & forms of information impact collective navigation"

#### A Additional Plots

In Figure S1 we show arrival time distributions in the *absence* of flow for the [FI] and [FA] models. We observe increased variance in the distribution for low levels of actual heading information. Similarly, in Figure S2 we show arrival time distributions for the [FI] and [FA] models in *unfavourable flow* ($u_y > 0$). Two sensing ranges are shown, namely $R = 20$ and $R = 500$. In Figure S3 we show arrival time distributions for the [FI] and [FA] models in a *real-world* flow. In Figure S4 we show analogous heatmaps to Figures 3 and 5, instead showing the navigational efficiency as defined in [1] and described in Equations (1),(2).Figure S4 shows that communicating intended headings is preferable in unfavourable flows. These plots align with our results in Section 3 using the median arrival time metric.

The remainder of the figures in this section are annotated versions of the heatmaps presented in Section 3. Labels in the heatmap cells give the median arrival time averaged over 50 realisations relative to solo navigators in a static environment. Figures 3 and 5 are annotated in Figure S5, Figure 6 in Figure S6 and finally, in Figure S7 we show the difference in median arrival time between the [FI] and [FA] models across a variety of flow strengths and sensing ranges in a real-world flow environment, analogously to Figure 3c).

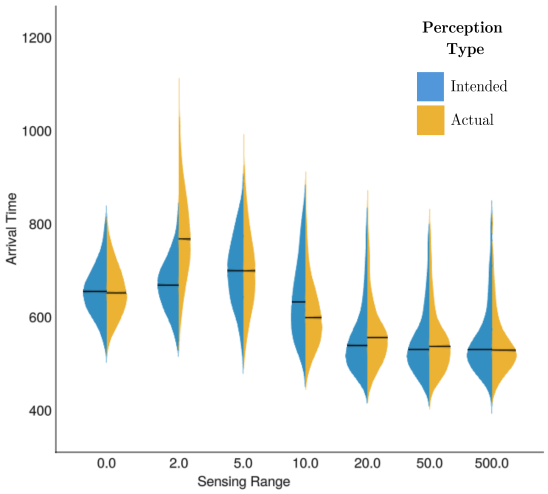


Figure S1: Arrival time distributions in the absence of flow for the [FI] (blue) and [FA] (yellow) models. Note the increased variance for low levels of actual heading information ($R \lesssim 10$). Black lines indicate the median arrival time. All distributions are averaged over 50 realisations.

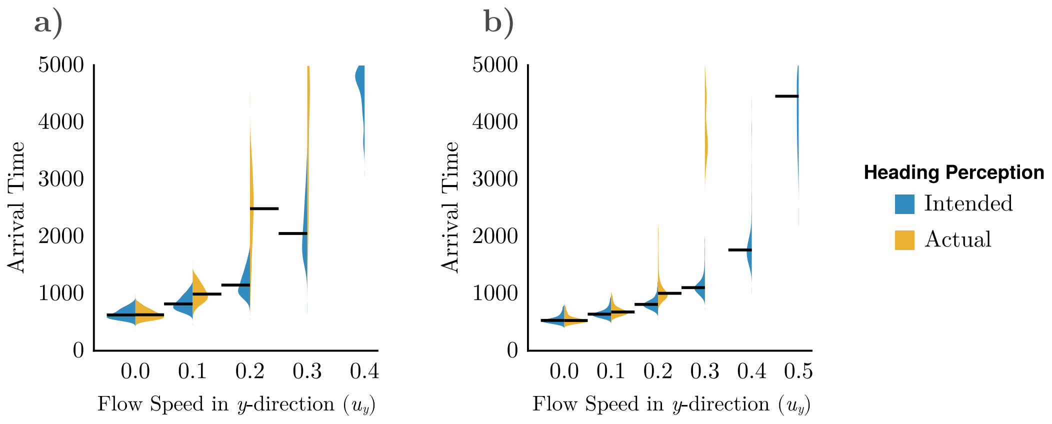


Figure S2: Arrival time distributions for the [FI] (blue) and [FA] (yellow) models in unfavourable flow ($u_y > 0$) for a) intermediate ($R = 20$) and b) extreme ($R = 500$) sensing ranges. Black lines indicate median arrival time – in the absence of a line, the migration failed. All distributions are averaged over 50 realisations.

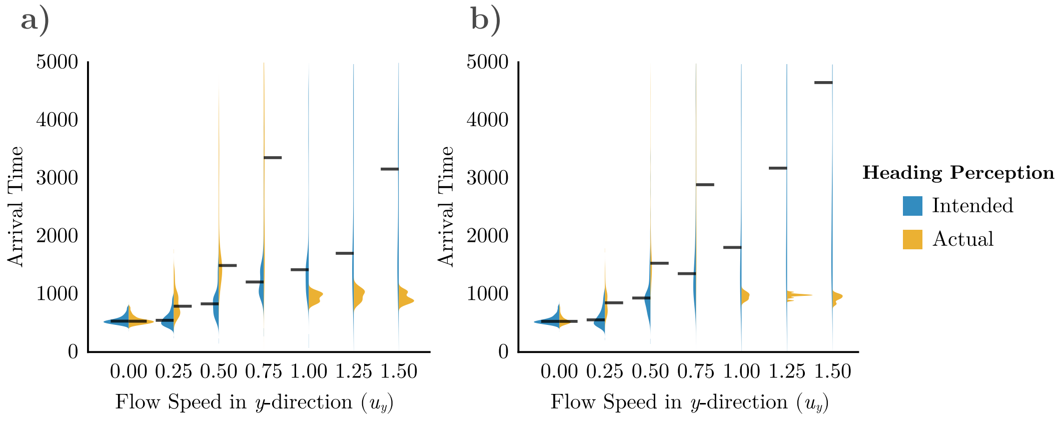


Figure S3: Arrival time distributions for the [FI] (blue) and [FA] (yellow) models in a real-world flow for a) $R = 50$, b) $R = 500$). Black lines indicate median arrival time – in the absence of a line, the migration failed.

### B Metric Sensitivity

It is necessary to define a metric of navigational success to compare model parameters/setups. To identify a robust and informative metric, we vary parameters in the initial configuration (distance to the goal, tolerance around the goal ($\varepsilon$) and the number of individuals) and assess the effect on a variety of metrics. Given the abstract modelling approach employed here, we seek fairly general measures that are relatively robust to parameter variation. We consider (i) the time of first arrival, (ii) the median arrival time, (iii) the time taken for 90% of the population to arrive, and (iv) the time of arrival of the last individual. We also consider the *navigational efficiency* of the group, based on discussions of this metric in [1, 2].

Figure S8 shows arrival time distributions across a variety of sensing ranges, and illustrates selected centiles of the distribution (Fig. S8a), the effect of the goal tolerance (Fig. S8b), and the effect of the size of the population (Fig. S8c). Even in the absence of flow, the distributions vary according to the sensing range [3]. The tails of the distributions, in particular, are poor representations of general migration success as they hide the trend apparent in the centres of the distribution. The tolerance around the goal has a large effect on the arrival distributions, especially when there is a high sensing range. This effect, however, largely manifests in the tails of the distribution, and has relatively little impact on the median arrival time. Determining the size of the goal is not the aim of the current study, hence a metric that is robust to changes in tolerance is desirable. Finally, we see that varying the number of agents has a sizeable effect on the arrival time distributions.

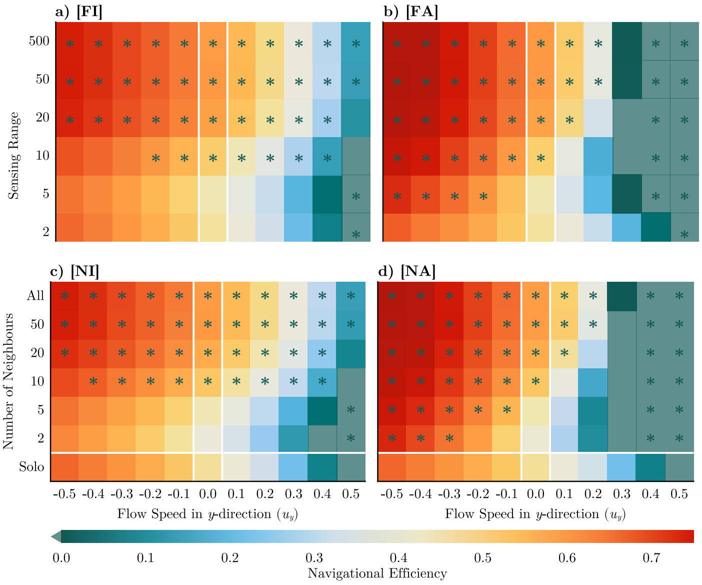


Figure S4: **Communicating intended headings gives higher navigational efficiency in opposing laminar flows.** (a-d) Heatmaps of the flow-affected navigational efficiency at $T = 250$ (Equation (2)) as a function of the flow speed in the $y$-direction and the sensing range. Each parameter combination is averaged over 50 realisations. a) Results for a fixed range interaction and intended headings [FI]. b) Fixed range interaction and actual headings [FA], c) Nearest neighbour interaction and intended headings [NI]. d) Nearest neighbour interaction and actual headings [NA]. Green asterisks denote parameters where the corresponding solo navigational efficiency was improved upon by $\geq 5\%$. Lighter green cells indicate where the efficiency was less than 0, i.e., the group on average moved away from the target.

This is expected under the ranged interaction as the effective group size (and thus the amount of information an individual receives from the group) is much lower when $N$ is reduced. Despite this, the pattern across sensing ranges is qualitatively similar. From this, we conclude that the median arrival time provides a suitably robust metric by which to measure the performance of the group. If less than 50% of individuals arrive within $T = 5000$, the group is said to have *failed*, and will be omitted from any figures. In all simulations we will use $N = 100$, as this ensures a group large enough to maintain a broad level of collective information, while also allowing relatively efficient simulation.

For further context, we also consider the *navigational efficiency* of a group. In

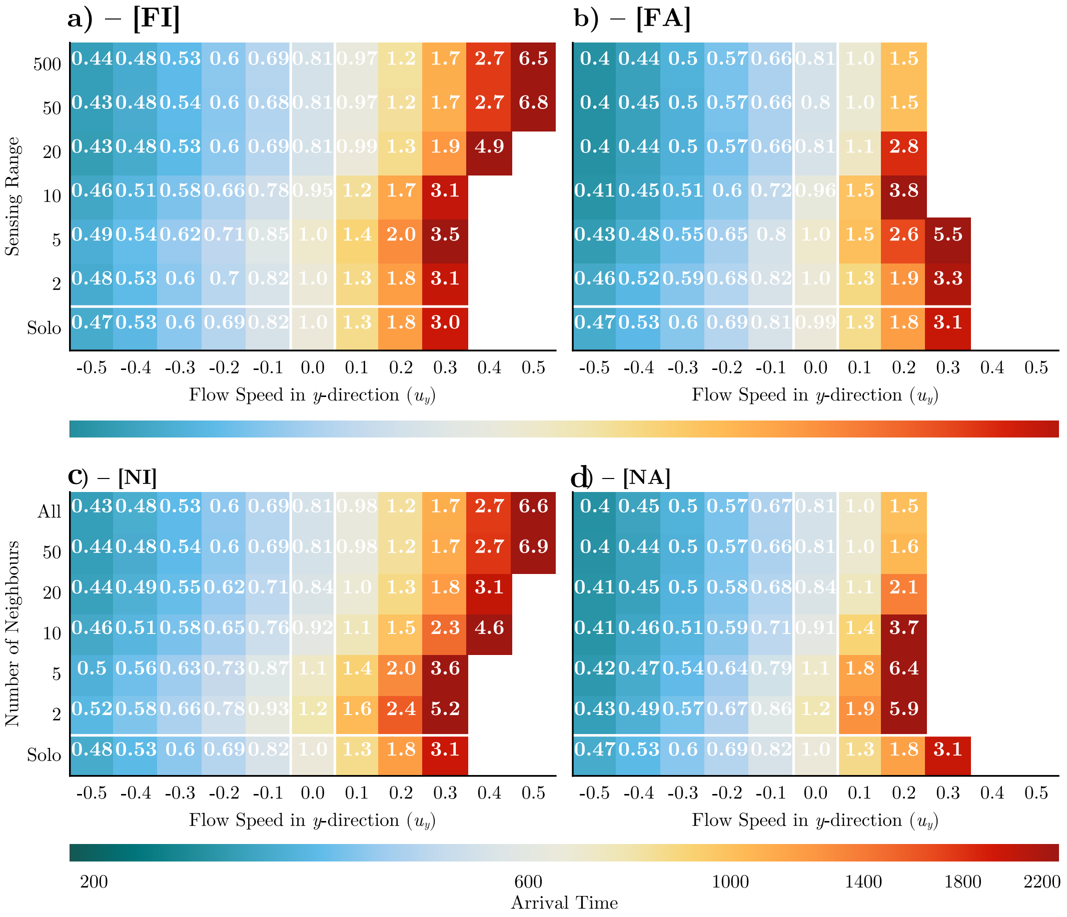


Figure S5: (a-b) Heatmaps of the median arrival time as a function of the flow strength and sensing range. Each parameter combination is averaged over 50 realisations. a) Results for a fixed range interaction and intended headings [FI] . b) fixed range interaction and actual headings [FA] . c) [NI] model. d) [NA] model. Annotations give the median arrival time, relative to a solo navigator in the absence of flow. Missing cells indicate where there is failed migration at the population level. All other parameters remain unchanged from Figures 3 and 5 of the main paper.

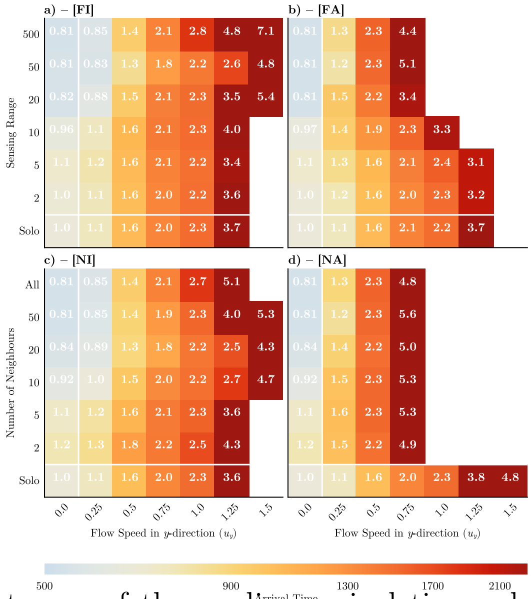


Figure S6: (a-d) Heatmaps of the median arrival time relative to an individual in the absence of flow, averaged over 50 realisations, for the [FI] , [FA] , [NI] and [NA] models respectively. All simulations are in a turbulent flow environment. Annotations give the median arrival time relative to a solo navigator in the absence of flow.

[1], this was defined as

$$E = \frac{d_0 - d_T}{T} \tag{1}$$

where $d_T$ is the distance from the centre of mass of the group to the centre of the target after $T$ timesteps of the simulation, and $d_0$ is the initial distance. In a laminar flow environment, we modify this to account for the negative (or positive) impact of the flow on individuals. For a constant flow in the $y$-direction with flow strength $\zeta$ and active swim speed $s = 1$, we define the *flow-affected navigational efficiency* as

$$\begin{aligned} E_f &= \frac{d_0 - d_T}{\|v + u\| T} \\ &= \frac{d_0 - d_T}{\|s \sin(-\pi/2) + \zeta \sin(\vartheta - \pi/2)\| T} && \text{considering only } y\text{-direction} \\ &= \frac{d_0 - d_T}{(s \pm \zeta) T} && \text{for } \vartheta = -\pi, 0 \text{ respectively.} \end{aligned} \tag{2}$$

Note that in the variable flow case, whether flow is beneficial or detrimental varies in time and position, and we thus omit efficiency plots for this case.

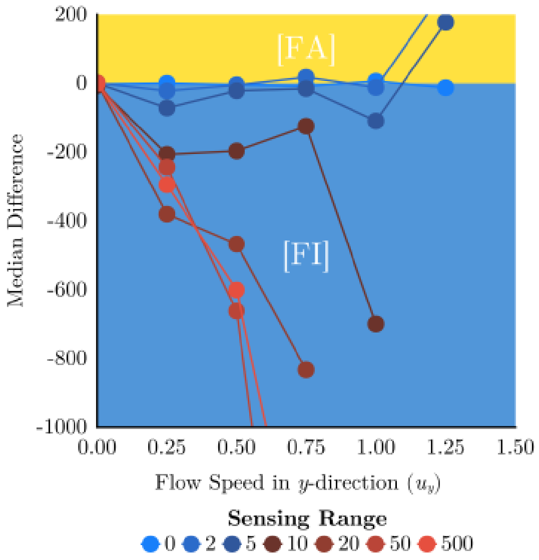


Figure S7: The difference in median arrival time between the [FI] and [FA] models across a variety of flow strengths and sensing ranges in a real-world flow environment. Positive values (yellow) indicate scenarios in which sharing actual headings leads to faster arrival times than when sharing intended headings (blue). Figure 6 shows heatmaps of this data for each model.

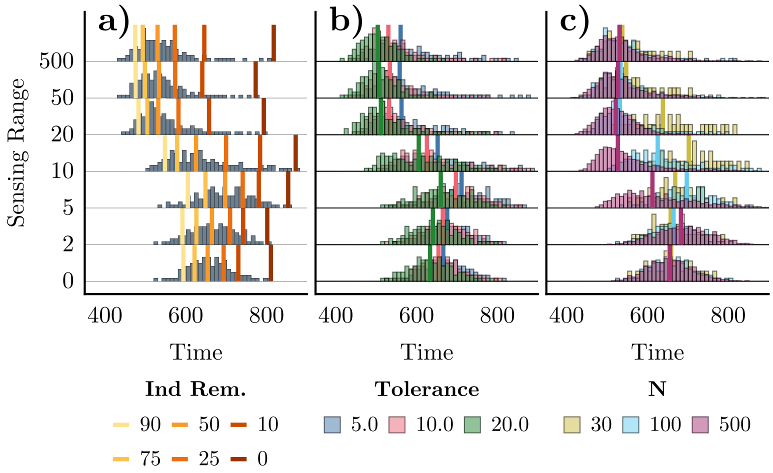


Figure S8: **Median Arrival Time is a Relatively Robust Metric.** In the absence of flow, the distribution of arrival times is shown across an average over 10 realisations. With 100 individuals and a goal tolerance of 10, the choice of metric affects the result: the final arrival time shows a very different pattern to more central centiles (a). However, varying the goal tolerance has little effect on the median arrival time despite the longer tails apparent for lower tolerances (b). As expected, changing the group size affects the median arrival time in this setup due to the smaller effective group size (c).

### C Sensitivity to Confidence in the Goal Direction

In this section we study the effect of changing the confidence in the goal direction, i.e., the concentration parameter of the inherent information distribution, $\kappa$, for both a laminar and a turbulent flow. Figure S9 largely reinforces intuition on the impact of $\kappa$. When the individual has more inherent information, arrival times are much lower. We still see a benefit to collective navigation when utilising intended headings, but this is only significant for strong unfavourable flows (Figure S9c). Collective navigation utilising actual headings provides a benefit in favourable flows, as seen previously, however continues to negatively impact on migration performance in unfavourable flows. If individuals have less information ($\kappa = 0.5$), collective navigation provides a benefit in all flows when using both intended and actual headings. In unfavourable flows, this effect is significant. This follows naturally from the observation that collective navigation is particularly beneficial when individuals have low confidence in the target direction according to their inherent information. In a turbulent flow setting, with low inherent information (Figures S9e-f) collective navigation provides a benefit for low flow strengths. However, as flow strength increases, the benefit weakens until the group begins to provide a negative impact on performance. Increasing the confidence in the goal direction in this setting (Figures S9g-h) shows a similar pattern, although the peak in benefit is much less pronounced due to the improved navigational capacity of the individual.

### D HYCOM Flow Data

HYCOM is a global ocean circulation model, validated against real-world data [4]. In the context of the present study, this ensures a realistic looking flow field, and we map the dimensional data of HYCOM onto our non-dimensional model domain. Note that the area chosen is from the North Atlantic between 13th April and 7th June 2021, shown in Figure 1d). This region intersects with the path of the Gulf Stream, therefore ensuring a region of strongly varying flow that intersects with the navigation route. The corners of the rectangle are at (34°N, 321.76°E), (34°N, 332.08°E), (59.76°N, 321.76°E) and (59.76°N, 332.08°E). All data are taken at surface depth (0 m). This region is then linearly mapped to the computational domain $(75, -75) \times (-50, 400)$ and normalised such that the mean flow strength (averaged over space and time) equates to the fixed flow strength parameter $\zeta > 0$. Note that if an individual leaves this region, the flow is wrapped periodically. Note further that HYCOM data can contain missing velocity values at certain points in time, which are mapped to zero prior to normalisation. HYCOM data was downloaded using the API provided by ERDDAP at `http://apdrc.soest.hawaii.edu/erddap/griddap/hawaii_soest_a2d2_f95d_0258`. This provides a NetCDF file, from which the missing values are removed before normalisation and the construction of a cubic spline interpolation function for each dimension (time, two-dimensional space and two-dimensional velocity). The interpolation function construction is provided by Interpolations.jl. Most missing values in the area chosen are due to the Azores Islands.

### E Calculating the Concentration Parameter

In Section 2, we described the use of a lookup table for estimating the concentration parameter $\kappa$. For a larger number of samples and larger $\kappa$, the maximum likelihood estimator proposed by Mardia and Jupp is usually used, see Algorithm S2. However, this was shown in [3] to be biased towards zero for low numbers of samples (neighbours in our case). We thus used a lookup table for estimation. This reduces the bias and smooths the distribution around $\kappa = 0$. Algorithm S1 shows a Julia implementation of calculating the circular resultant vector, and Algorithm S2 shows how this is used to estimate $\kappa$. Finally, Algorithm S3 describes how the lookup table is used if $\kappa$ is small.

```
function circ_resultant(
    samples::Vector{Float64},
    weights::Vector{Float64}
)::Float64
    R = sum(weights .* exp.(im .* samples))
    R = abs(R) ./ sum(weights)
    return R
end
```

Algorithm S1: Calculation of the circular resultant vector.

```julia
function mardia_jupp_kappa_mle(
    headings::Vector{Float64},
    weights::Vector{Float64}
)::Float64
    headings = headings[:]
    N = length(headings)
    if N > 1
        R = circ_resultant(headings, weights)
    else
        R = headings
    end

    if 0 <= R < 0.53
        κ = 2 * R + R^3 + 5 * (R^5) / 6
    elseif 0.53 <= R < 0.85
        κ = -0.4 + 1.39 * R + 0.43 / (1 - R)
    else
        κ = 1 / (R^3 - 4 * R^2 + 3 * R)
    end
    return κ
end
```

Algorithm S2: Mardia-Jupp method for calculating the maximum likelihood estimator of $\kappa$.

```
function get_kappa(
    headings::Vector{Float64},
    weights::Vector{Float64},
    kappa_CDF,
    kappa_input,
)::Float64
    κ = mardia_jupp_kappa_mle(headings, weights)
    N = length(headings)
    if κ < 25 && N < 25
        kappa_lookup_index = round(Int64, κ * 20) + 1 # equivalent
         ↪  to line below
        # _, kappa_lookup_index = findmin(abs.(κ - kappa_input))
        cdf_sample = rand()
        temp = findfirst(x -> cdf_sample < x, kappa_CDF[:, N-1,
         ↪  kappa_lookup_index])
        κ = kappa_input[temp] + rand() * (kappa_input[2] -
         ↪  kappa_input[1])
    end
    return kappa
end
```

Algorithm S3: Lookup table for small values of $\kappa$ and neighbours. `kappa_cdf` and `kappa_input` are available online at `https://melbourne.figshare.com/articles/dataset/kappaCDFLookupTable_mat/14551614` as a `.MAT` file. To improve performance in the present study, this was converted to a Julia-native `.jld2` file.

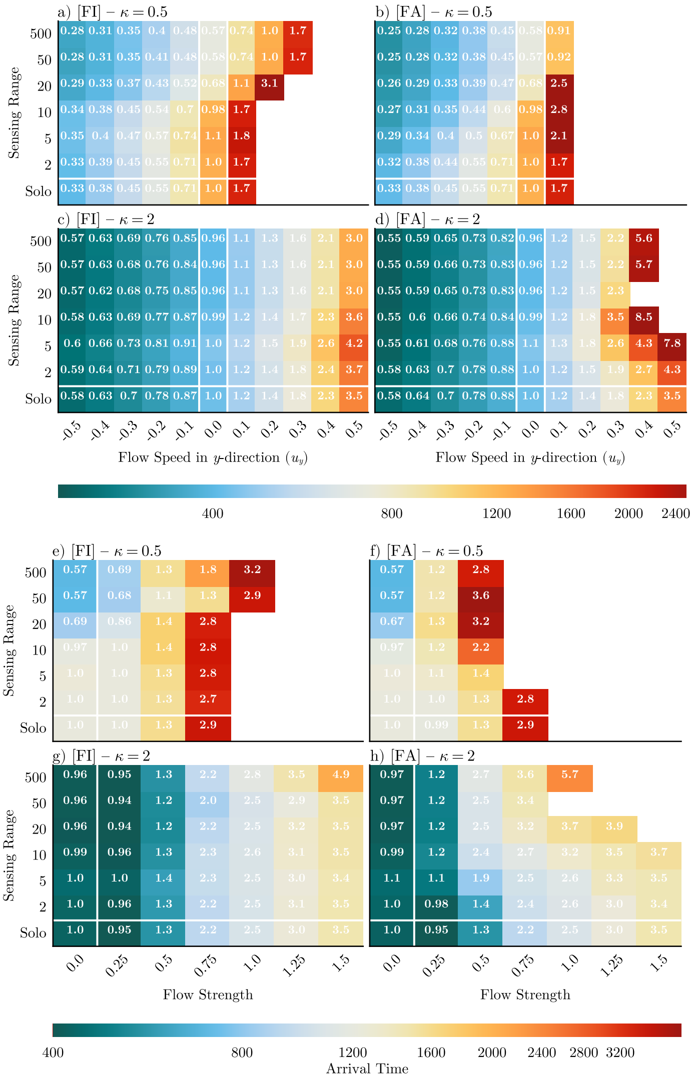

Figure S9: **Varying the confidence in the goal direction affects efficacy of collective navigation.** (a-d) In the case of laminar flows, all parameters are the same as in Figure 3 except for the concentration parameter of the inherent information distribution $\kappa$. (a-b) show the [FI] and [FA] models, respectively, with $\kappa = 0.5$ while (c-d) show the case when $\kappa = 2$. (e-h) In the case of turbulent flows, all parameters are the same as in Figure 6 except for the concentration parameter of the inherent information distribution $\kappa$. (e-f) show the [FI] and [FA] models, respectively, with $\kappa = 0.5$ while (g-h) show the case when $\kappa = 2$. Note the concentration parameter is lower ($\kappa = 1$) in all other figures. Annotations in all panels gives the median arrival time relative to a solo navigator in the absence of flow.
